# Late cortical dynamics mediate the symmetry-induced numerosity illusion

**DOI:** 10.64898/2026.02.27.708506

**Authors:** Alessandro Benedetto, Roberto Arrighi, Elisa Castaldi

## Abstract

Numerosity perception enables us to quickly estimate the number of objects in a visual scene, a fundamental skill supporting efficient interaction with the environment. In this study, we tracked how the human brain transforms physical inputs into numerical percepts. Using EEG we measured the neural dynamics to assess whether numerosity is perceived directly or inferred from correlated non-numerical features, and how these signals are integrated with contextual cues that bias perception. We leveraged on a visual illusion in which dot arrays appear less numerous when arranged symmetrically rather than randomly. Participants viewed dot arrays of varying physical or perceived numerosity, illusorily altered by the dots’ spatial arrangement. Both univariate and multivariate analyses revealed that physical numerosity was decodable from occipitoparietal electrodes as early as ∼50 ms after stimulus onset, independently of low-level features such as dot size or convex hull. Spatial arrangement was represented later, from ∼150 ms, and perceived numerosity, biased by symmetry-induced grouping, emerged at a similar latency. These results indicate that early neural signals encode physical numerosity directly, whereas the symmetry-induced underestimation arises from later grouping processes that integrate spatial information to shape perceived numerosity.

## Introduction

The human visual system is capable of quickly and accurately perceiving numerosity, that is the number of objects or individuals in a visual scene. This ability is present early in life and is shared across many species^1–10^, suggesting an evolutionarily conserved “number sense” that provides key ecological advantages such as guiding decisions about threat detection, foraging, and social interactions. Consistent with its relevance for survival, converging evidence shows that numerosity discrimination is both fast and automatic. Humans, for example, can provide numerosity-based oculomotor responses with saccadic latencies as short as 190 milliseconds^11^, and even the pupillary light reflex – one of the most automatic physiological responses – is modulated by numerosity^12,13^. Consistent with this automaticity, both humans and non-human primates base their choices in tasks such as odd-one-out discrimination on numerosity rather than other task-relevant visual attributes like area or density^14,15^. Moreover, in Stroop-like paradigms, numerosity can bias the perception of other quantitative dimensions^16^, further underscoring its perceptual dominance. Numerosity perception adheres to Weber’s law and exhibits adaptation effects, leading several authors to propose that number represents a fundamental visual attribute directly encoded by dedicated neural mechanisms^17–19^. Neurophysiological findings further bolster this view, revealing neurons selectively tuned to numerosity across species—from monkey^3,4,9^ and crows^20,21^ to humans^22^. Yet, despite evidence for specialized mechanisms, the precise neural computations supporting this capacity remain elusive.

First, it is still debated whether number is sensed directly or inferred from correlated visual features—such as area, density, or item size—since variations in numerosity inevitably co-occur with changes in these non-numerical dimensions. Previous EEG and fMRI studies in humans have investigated how non-numerical properties of numerical stimuli affect brain activity^23–29^. These studies found that numerosity-related signals emerge rapidly—within about 75 ms—and originate from early visual regions, even after controlling for certain non-numerical factors. However, most analyses examined only one non-numerical variable at a time or relied on mathematically defined constructs of size and spacing that do not directly correspond to perceptual features^30^. Only two fMRI studies have recently introduced a method better capable of disentangling neural responses to numerical and non-numerical features, overcoming the limitations of earlier work^31,32^. This method involves orthogonally varying perceptually meaningful dimensions and simultaneously accounting for all of them using multiple regression representational similarity analysis (RSA)^33,34^. The results revealed that numerosity-specific representations are present starting from primary visual areas and extending across both the dorsal and ventral pathways^31,32^. However, the limited temporal resolution of fMRI makes it impossible to determine whether these responses reflect feedforward encoding of numerosity or arise from later feedback processes that compute number indirectly. As a result, from these studies it remains unclear whether fast neural responses reflect a dedicated mechanism that encodes numerosity directly or through the combination of multiple correlated non-numerical features.

Second, even assuming that numerosity-specific signals are present, it remains largely unknown how these signals are integrated with contextual cues that can bias our perception of number. Indeed, although numerosity perception is largely automatic, it is strongly influenced by spatial organizational cues^35–38^. For instance, when items are linked by thin lines, the perceived numerosity of an array is systematically reduced^39–41^. This effect, also known as the connectedness illusion, is also observed when connections are only implied (such as in Pacman-like stimuli) or when items are enclosed within shapes (such as ovals), suggesting that physical linking is not necessary to elicit the underestimation illusion^41–43^.

Another influential spatial organizational cue that shapes numerosity perception is symmetry, a feature with important biological relevance that supports object identification and figure-ground segregation. For example, the symmetric patterns of the lionesses’ eyes and stripes on their fur allow us to readily segment their three camouflaged heads from the grass as they prepare to hunt, rather than perceiving six unrelated black spots and lines. In laboratory settings, a seminal study demonstrated that symmetrical dot patterns are consistently perceived as less numerous than randomly arranged arrays^44^. Early explanations of this phenomenon suggested that the underestimation arises because symmetry introduces redundancy (input autocorrelation), prompting observers to attend to only part of the display and thereby underestimate the total number of elements^44^. We recently proposed an alternative explanation for the symmetry-induced numerosity underestimation, suggesting that it arises from perceptual grouping processes that attempt to support the system in identifying perceptual units^45,46^. Symmetric elements may be perceptually grouped into unified shapes, which in turn decreases the perceived numerosity of the array. In line with this possibility we previously demonstrated that the symmetry-induced underestimation is more pronounced for moderate numerosities typically processed by the Approximate Number System (ANS), and diminishes in high-density arrays, where perception relies more on texture-based mechanisms and object segmentation is more difficult^45,47,19^. Furthermore, the degree of numerosity underestimation grows as the grouping cue becomes stronger, with additional axes of symmetry producing greater reductions in perceived number. We also found that individual differences in perceptual style modulate the symmetry-induced underestimation effect: individuals with higher autistic traits, who tend to adopt a more local perceptual style, show reduced underestimation. Finally, diverting attention to a concurrent color–orientation conjunction task significantly reduces the symmetry-induced numerosity underestimation, suggesting that this effect may depend on the distribution of attention across elements as they are bound into a single perceptual object^46^. Together, these findings suggest that the visual system estimates numerosity from perceptually segregated units or objects defined by grouping cues, rather than from individual elements. Despite strong behavioral evidence supporting this view, how the brain constructs a representation of perceived numerosity by integrating physical inputs with grouping cues remains largely unexplored.

The current study has two goals. First, it aims at determining whether early neural responses directly encode the numerosity of the dot array, or whether they arise from integrated responses to non-numerical features from which numerosity can be inferred. For the first time, we applied multiple regression representational similarity analysis (RSA), a method previously used only in fMRI studies, to neural responses at each time point, enabling us to simultaneously account for perceptually defined non-numerical features and isolate the variance uniquely attributable to numerosity across the time course. Secondly, assuming that early signals genuinely reflect physical numerosity, we sought to track when and how the brain integrates these signals with spatial configurational cues to construct perceived numerosity. We hypothesized that if symmetry reduces perceived numerosity by binding symmetric elements into unified shapes, its neural signature should emerge at relatively late processing stages, reflecting the activation of grouping mechanisms that support object segmentation. Indeed, this would be in line with the report of a study about connectedness. Fornaciai and Park^27^ demonstrated that in the processing of connected items – usually perceived as reduced in numerosity relative to a set of independent elements – the neural activity occurring in area V2 before 100 ms after stimulus onset reflects physical numerosity while the signature of connectedness biasing perceived numerosity emerges later, at around 150 ms and in a different area of visual hierarchy, namely V3). To anticipate the results, we found that that numerosity is represented in multiple stages: an early stage reflecting physical numerosity and a later stage corresponding to perceived numerosity, both independently of low-level visual features. The later stage interacts with symmetry-based grouping cues, suggesting that the symmetry-induced underestimation arises from later-stage perceptual grouping mechanisms.

## Results

### Sequential encoding of numerosity and symmetry revealed by univariate VEP dynamics

Participants viewed a series of centrally presented dot arrays, of various numerosity arranged either symmetrically or randomly, while continuous EEG recordings were acquired (Figure 1). On occasional trials (10%), indicated by a color change to red of the stimuli, participants were asked to compare the numerosity of the current array to the previous one and respond by pressing one of two arrow keys to indicate whether the current array contained more or fewer dots than the previous. This task was included to ensure that participants maintained attention to the stimuli; however, these catch trials were excluded from all subsequent analyses.

**Figure 1.**
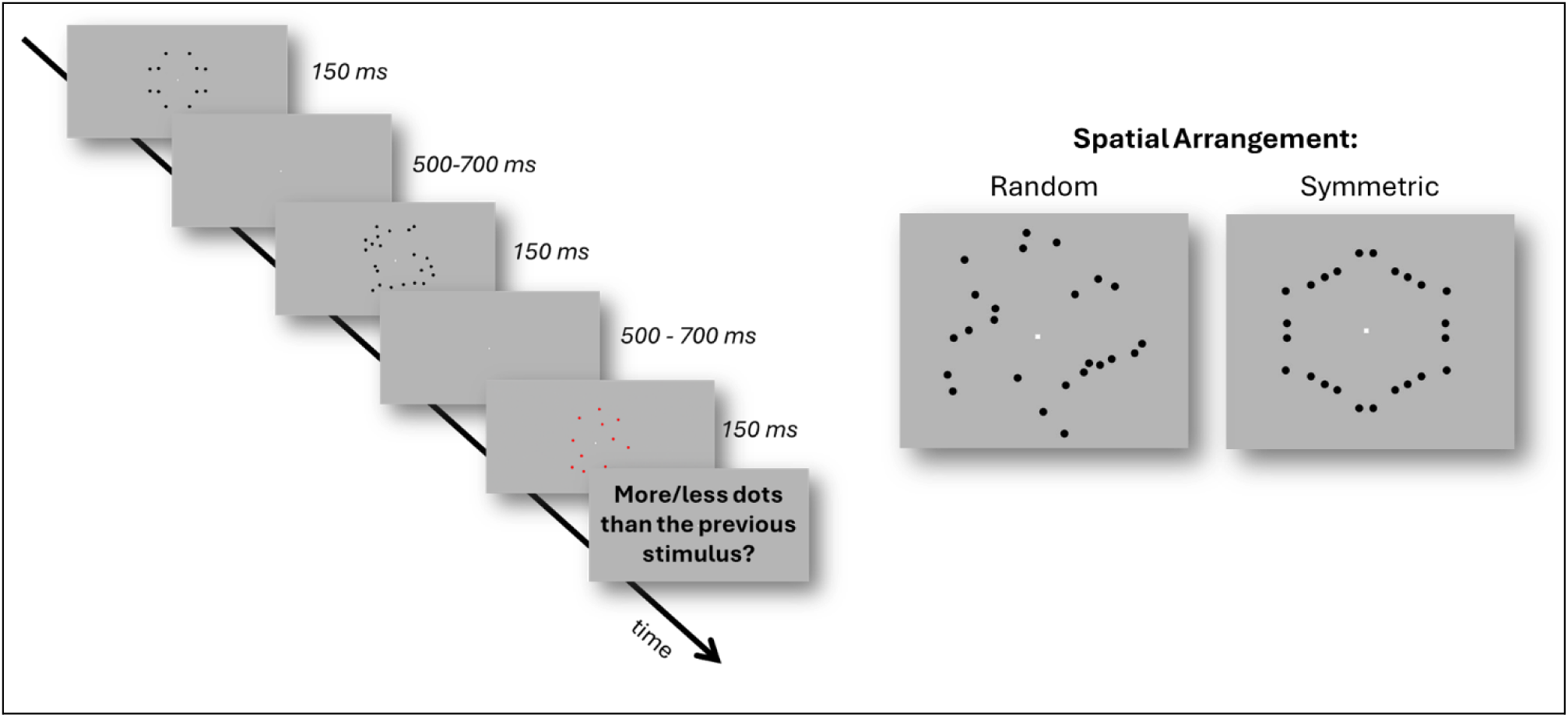
Experimental design and stimulus examples. Participants viewed centrally presented dot arrays of varying numerosity, arranged either randomly or symmetrically. On occasional catch trials, the array color changed to red, prompting participants to compare the current numerosity with that of the preceding array and respond via arrow keys (“more” or “less”). The left panel illustrates a sequence of trials, including a red catch trial, while the right panel shows example stimuli for the random and symmetric conditions.

To provide an initial overview of the effects of numerosity (12 vs. 24) and spatial arrangement (symmetry vs random), we first examined the topographical distribution of brain activity across the entire scalp, without restricting the analysis to any ROI (Figure 2A). The results revealed the emergence of two distinct spatiotemporal clusters associated with the main effects of numerosity and spatial arrangement (p < 0.01). As shown in the top row of Figure 2A, VEPs to different numerosities (12 vs. 24) could be distinguished at early latencies, whereas the main effect of spatial arrangement emerged later in the time course (bottom row of Figure 2A). Both effects were detected over occipito-parietal electrodes, which we selected as ROI, and their time course is shown in Figure 2B. A significant effect of numerosity emerged at two distinct time windows: an early and a later phase. The early numerosity effect appeared as soon as ∼80 ms post-stimulus, remaining significant until approximately 100 ms. During this interval, VEPs elicited by 12-dot arrays were more positive than those elicited by 24-dot arrays, largely independent of spatial arrangement. A second numerosity effect emerged around 170 ms, persisting until about 200 ms. In this later window, brain responses were again modulated by numerosity (this time with more positive amplitudes for 24 than 12) and also by spatial arrangement, with random arrays evoking more positive responses than symmetric ones. The main effect of symmetry appeared slightly earlier than the second numerosity window yet still emerging at a relatively late time range, beginning around 150 ms and remaining sustained up to 470 ms post-stimulus.

**Figure 2.**
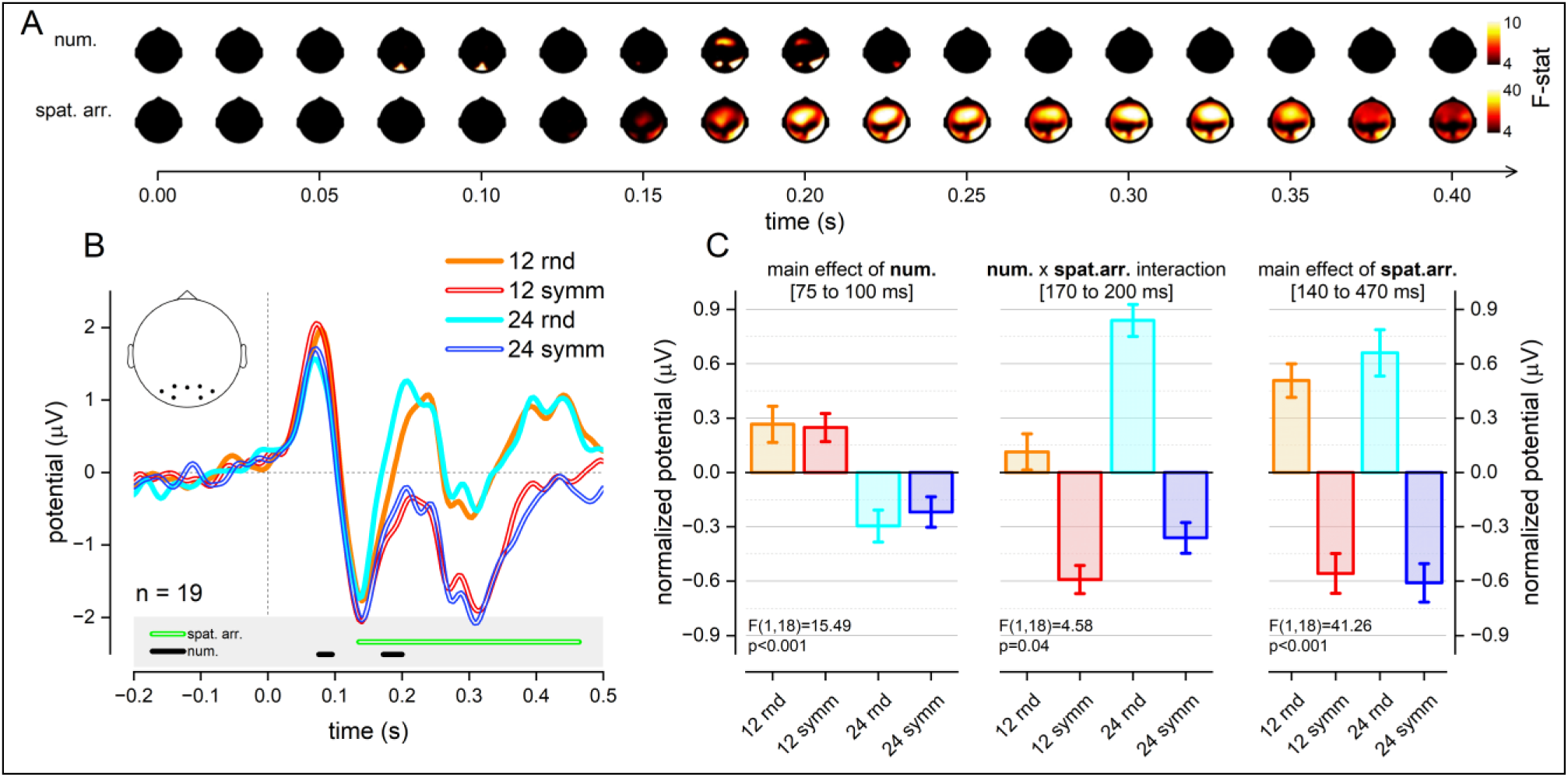
Results of the linear mixed-effects model on VEPs. (A) Topographic maps of F-statistics highlighting electrodes most responsive to the main effects of numerosity (top row) and symmetry (bottom row). (B) Grand-averaged VEP waveforms from ROI occipitoparietal electrodes (inset). (C) Statistical results from the region-of-interest analyses for the three-time windows of interest (75–100 ms, 170–200 ms, and 140–470 ms), illustrating the emergence of an early numerosity-specific component, followed by a later interaction between numerosity and symmetry, and a sustained main effect of symmetry.

To estimate the magnitude of the effects and characterize them more precisely, we extracted three representative time windows within the occipitoparietal ROI based on the temporal profile of the univariate results (Figure 2C). Activity in the very early latency window (75–100 ms) showed a main effect of numerosity (F(1,18) =15.49, p < 0.001), but no effect of symmetry (F(1,18) = 0.10, p = 0.735) or the interaction (F(1,18) = 0.44, p = 0.515). This indicates that early cortical responses are selectively sensitive to physical numerosity, independent of spatial arrangement. In a later time window (170-200 ms), both numerosity and spatial arrangement produced significant main effects (F(1,18) = [14.99, 31.44], p = [0.001, <0.001], respectively), and their interaction was also significant (F(1,18) = 4.58, p = 0.046), suggesting that symmetry begins to modulate VEP responses (with such modulation depending on numerosity) only after the initial stage in which the signals did not reflect spatial arrangement. This is consistent with prior reports showing symmetry-induced underestimation effects of approximately 12% and 14% for 12 and 24 dots, respectively^45^. Finally, analyses within the extended window corresponding to the sustained symmetry-related activity (140-470 ms), revealed a strong main effect of symmetry (F(1,18) = 41.26, p < 0.001), with no effect of numerosity (F(1,18) = 0.30, p = 0.585) or interaction (F(1,18) = 1.56, p = 0.226).

### Disentangling the contributions of numerical and non-numerical dimensions using multiple regression RSA

To determine whether EEG modulations were specifically driven by numerosity rather than by non-numerical features such as item size or convex hull, we applied Representational Similarity Analysis (RSA). RSA allows the simultaneous assessment of multiple dimensions, both numerical and non-numerical, in relation to evoked responses.

For each time point, a neural representational dissimilarity matrix (neural RDM) was constructed by computing the correlation distance between response patterns across all condition pairs. This matrix quantifies the degree of dissimilarity between VEPs elicited by different stimuli. Neural RDMs were then entered into a multiple regression analysis with theoretical representational dissimilarity matrices (theoretical RDMs) as predictors. Each predictor captured dissimilarities along a specific stimulus property, including numerosity, spatial arrangement, item size, and convex hull. Importantly, numerosity, item size, and convex hull were orthogonally manipulated within the stimulus space, ensuring that the corresponding predictors were decorrelated.

This approach yielded beta weights for each predictor at every time point, reflecting the variance in neural dissimilarity uniquely explained by each stimulus dimension, above and beyond the contribution of all the others. Consequently, this method allowed us to evaluate the extent to which VEP differences could be attributed to numerosity independently of other quantitative dimensions or characteristics of the stimuli.

Figure 3 depicts the time-resolved beta weights for each predictor. Beta weights different from zero (as indicated by the continuous thick lines) indicate that information related to that stimulus dimension contributes to the VEPs, over and above the contribution of the other dimensions. Beta weights for numerosity (black line), revealed two distinct significant time windows: an early window spanning 70–100 ms and a later window from 160–220 ms. Beta weights for convex hull (red line) were significant between 60 and 280 ms, whereas item size (blue line) did not produce significant effects at any time point. Beta weights for spatial arrangement (green line) became significant no earlier than 160 ms and persisted until 450 ms. Overall, these results indicate that VEPs are modulated by numerosity independently of other visual stimulus features, supporting the notion of an early and sustained neural sensitivity to numerical information.

**Figure 3.**
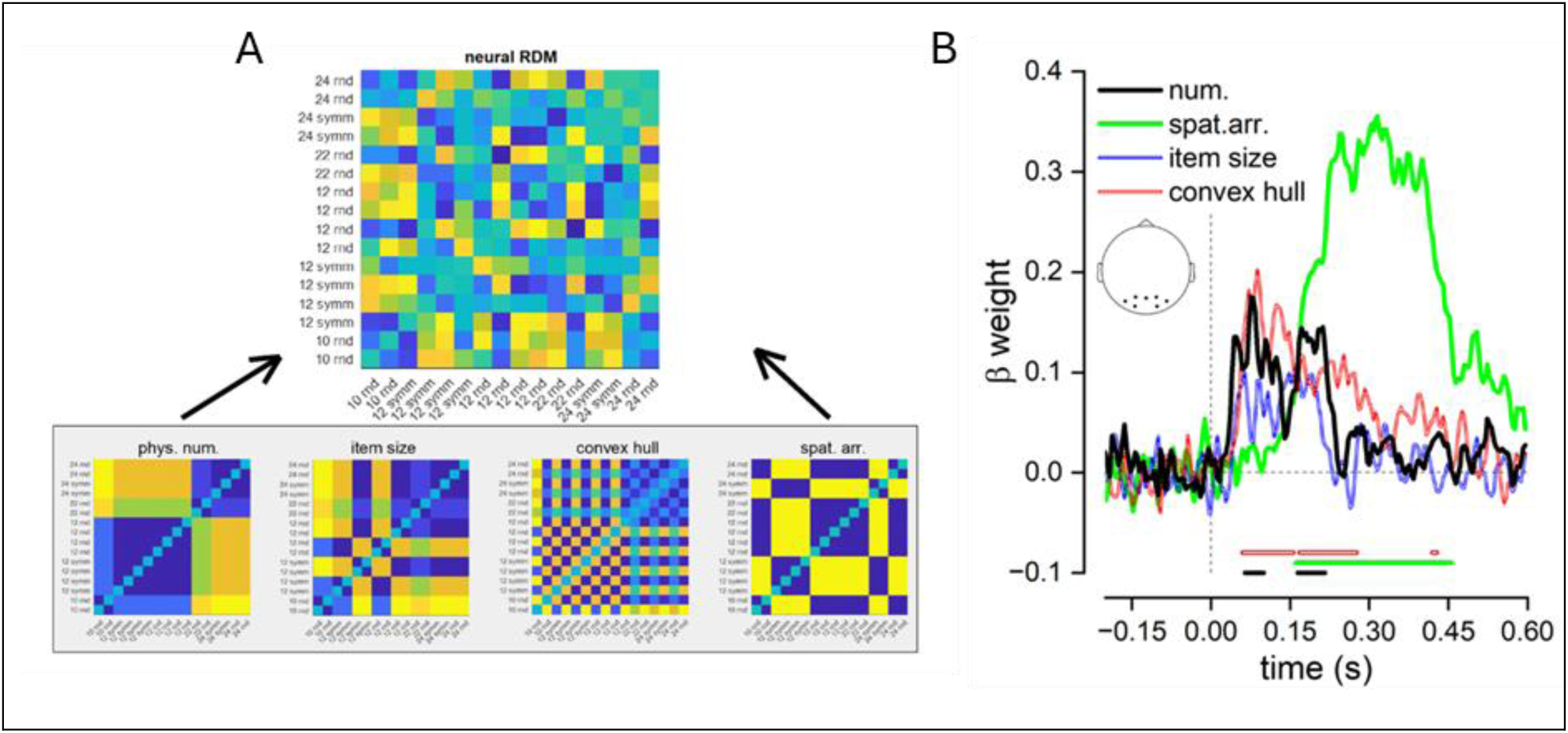
Representational Similarity Analysis (RSA) of EEG responses. A, top: Neural representational dissimilarity matrix (RDM) showing the pairwise correlation distances between VEP patterns for all stimulus conditions. A, bottom: Theoretical RDMs for each stimulus dimension: numerosity (phys. num.), item size, convex hull, and spatial arrangement (spat. arr.). Each matrix reflects predicted dissimilarities between conditions based on that specific stimulus property. B: Time-resolved beta weights from multiple regression RSA, representing the unique contribution of each stimulus dimension to neural dissimilarity over time. Shaded lines indicate the β weights for numerosity (black), spatial arrangement (green), item size (blue), and convex hull (red). Horizontal bars indicate time periods during which beta weights were significantly greater than zero (FDR corrected). The results demonstrate that VEP patterns are modulated early by numerosity and convex hull, whereas spatial arrangement effects emerge later.

### Disentangling perceived versus physical numerosity with multiple regression RSA

We next examined whether the symmetry-induced underestimation of numerosity could be detected using RSA. To this end, beta weights derived from multiple regression RSA were compared across two models incorporating numerosity, item size, spatial arrangement, and convex hull. In one model, the numerosity RDM reflected physical numerosities, whereas in the other, it reflected perceived numerosities, with lower values assigned to symmetrically arranged patterns to capture the symmetry-induced underestimation of numerosity. Specifically, arrays of 12 symmetric dots were modeled as containing 10 dots, and those with 24 symmetric dots as 22, based on the expected underestimation effect derived from average previous behavioral results^45^. Figure 4 depicts beta weights for physical numerosity (black) and perceived numerosity (gray) over time. In a relatively late time window (170-240 ms), neural RDM was significantly better accounted for by the perceived numerosity RDM than by the physical numerosity RDM (p < 0.05, FDR corrected). Notably, this effect appeared to overlap almost perfectly with the interaction window observed in the univariate analysis (Figure 1C), which showed an interaction between numerosity and spatial arrangement between 170 and 200 ms. Focusing on this same temporal window, we directly compared beta weights for perceived and physical numerosity. This analysis revealed a highly consistent effect across participants, with 16 of 19 showing higher beta weights for the perceived RDM compared to the physical RDM (t(18) = 3.75, p = 0.001. See inset in Figure 4). Together, these results indicate that VEP modulations in this time window are more closely predicted by perceived than physical numerosities, likely reflecting the symmetry-induced underestimation of numerosity.

**Figure 4.**
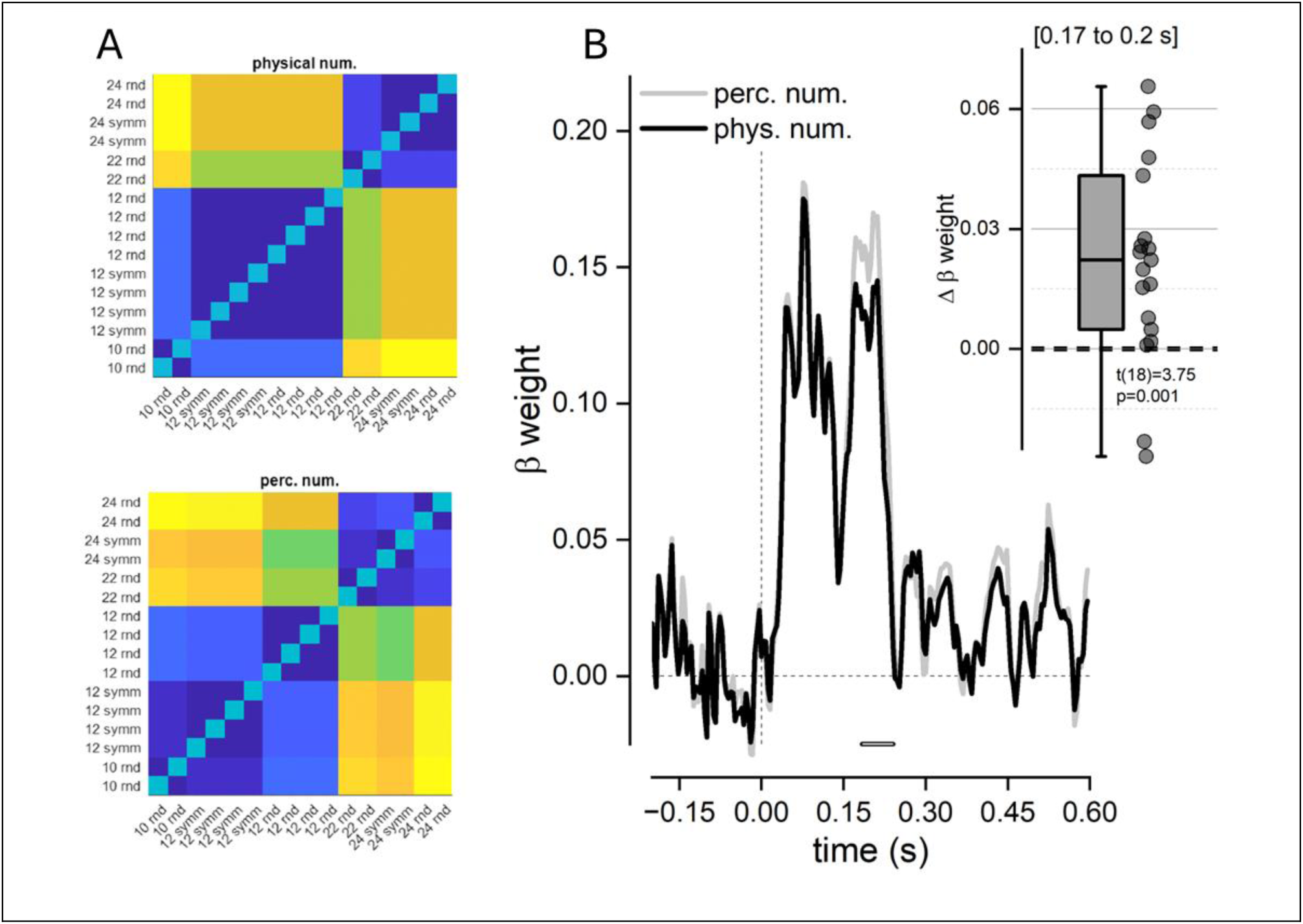
Neural representations of physical versus perceived numerosity. A: Representational dissimilarity matrices (RDMs) based on physical numerosity (phys. num., left) and perceived numerosity (perc. num., right) for all stimulus patterns. Color scale reflects dissimilarity values, with higher values indicating greater dissimilarity. B: Time-resolved beta weights from multiple regression RSA comparing the explanatory power of perceived (gray) versus physical (black) numerosity RDMs on neural data. Inset: Boxplot showing the difference in beta weights (Δβ) between perceived and physical numerosity in the 170–200 ms time window; most participants show higher beta weights for perceived numerosity (t(18) = 3.75, p = 0.001). These results indicate that neural activity in this late time window is more closely predicted by perceived than by physical numerosity, reflecting the symmetry-induced underestimation of numerosity.

### Sustained versus transient neural representations of numerosity and spatial arrangement revealed by MVPA

To examine the temporal dynamics of neural representations, we next performed a multivariate decoding analysis across all EEG channels. At each time-point, an LDA classifier was trained on the spatial pattern of EEG activity evoked by different numerosities or spatial arrangements. Specifically, classifiers were trained to distinguish between 12– and 24-dot stimuli and were subsequently tested on independent trials not used for training. Classification accuracy significantly above chance (50%) indicated that neural activity patterns were sufficiently distinct to generalize beyond the training set. An analogous procedure was used to decode spatial arrangement, with classifiers trained and tested on trials contrasting random versus symmetric configurations.

Figure 5A shows classification accuracies for discriminating numerosities and spatial arrangements at corresponding time points. Consistent with VEP and RSA results, numerosity could be decoded significantly above chance beginning at about 50 ms, peaking around 100 ms, and remaining significant until 350 ms, with a brief resurgence from 450 to 485 ms. In contrast, decoding of spatial arrangements emerged later, at about 115 ms, and remained significantly above chance for nearly the entire time course, reaching a peak of accuracy at about 230 ms. Overall, classification accuracy was substantially higher for spatial arrangement than for numerosity, indicating that the neural patterns associated with different spatial configurations were more distinct and thus more easily separable than those associated with different numerosities.

**Figure 5.**
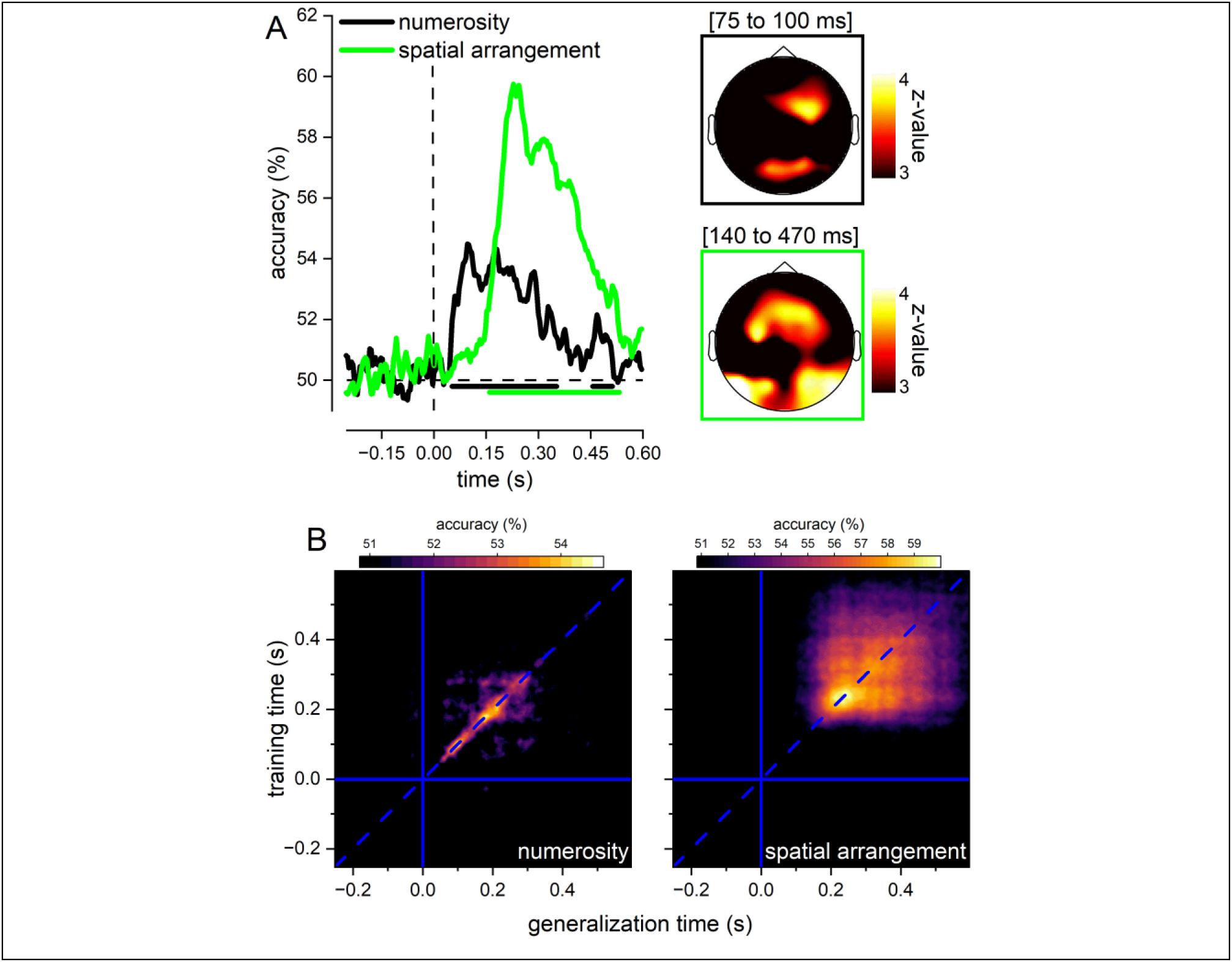
Multivariate neural decoding of numerosity and spatial arrangement. (A) Time-resolved classification accuracies, reveal two distinct peaks: an early numerosity-related effect around 100 ms and a later symmetry-related effect around 200 ms. These peaks highlight a temporal dissociation in the brain’s processing of quantity and spatial arrangement. Colored bars beneath the graph indicate statistically significant time intervals for each condition (black: numerosity; green: spatial arrangement). (B) Temporal generalization matrices show how well the neural representations for numerosity, and spatial arrangement generalize across time, with classifiers trained at a specific time point tested across all others.

Searchlight analysis revealed that classification of numerosity was primarily driven by activity in occipitoparietal and frontal channels, whereas discrimination of spatial arrangements was predominantly supported by occipitoparietal and central channels (Figure 5B-C).

We next assessed whether neural activity patterns generalized across time using temporal generalization analysis, which evaluates the extent to which information encoded at one time point can be reliably decoded at other time points, providing insights into the recurrence and temporal persistence of neural representations^48^. Figure 5D presents the temporal generalization matrices for numerosity (left) and spatial arrangement (right). For numerosity, classification accuracy was largely confined along the diagonal of the matrix, particularly at earlier latencies, indicating that the classifier performed best when trained and tested on the same time points. This pattern suggests that early numerosity-related neural responses are transient and time-specific. At later latencies, the generalization pattern became slightly more dispersed, suggesting a transition toward a more sustained processing. In contrast, classification of spatial arrangement was more temporally distributed. Above-chance cross-temporal decoding extended over a broader range, with reliable generalization observed up to about 200 ms, indicating that information about spatial configuration was encoded in more stable and enduring neural patterns.

## Discussion

In this study, we investigated how the brain builds a representation of perceived numerosity, starting from physical inputs and integrating the spatial arrangement of individual elements. Participants viewed a series of centrally presented dot arrays of varying numerosities, arranged either symmetrically or randomly, while continuous EEG recordings were collected. The results revealed that specific numerosity-related neural activity could be dissociated into two distinct temporal phases: an early phase (starting from ∼50-80 ms post stimulus) reflecting the encoding of “raw” numerosity, and a later phase (starting from ∼170 ms post stimulus) corresponding to perceived numerosity, which is modulated by segmentation processes prompted by symmetry, a property itself represented only relatively late in the timecourse.

We applied multiple regression representational similarity analysis (RSA) to EEG signals to isolate the variance uniquely attributable to numerosity while simultaneously account for perceptually defined non-numerical quantities and observed an activity related to numerosity emerging very early, beginning around 50 ms after stimulus onset. This activity was specifically bounded to numerosity over and above other non-numerical features such as item size, spatial arrangement, and convex hull. This finding aligns with a recent fMRI-MEG fusion study employing the same analysis method, which reported that numerosity-specific signals appear early (approximately 75 ms after stimulus onset), preceding the neural encoding of pairs of non-numerical features (e.g., total surface area and average item area) from which numerosity could be indirectly inferred^49^. These results support the notion that numerosity is encoded directly, rather than being derived from combinations of non-numerical cues.

Crucially, this initial representation of numerosity may then be revised by interactions with segmentation processes such as symmetry that define perceptual units. This emerged clearly when we compared beta weights between two models: one incorporating physical numerosity in the representational dissimilarity matrices (RDMs), and the other modeling perceived numerosity, with lower values assigned to symmetrically arranged patterns to capture the underestimation effect. Consistent with the idea that segmentation processes revise the initial numerosity representation, in a later temporal window (170–240 ms), neural RDMs were better accounted for by the perceived numerosity model than by the physical numerosity model, suggesting that perceived numerosity is represented after symmetry-based grouping.

To further investigate the temporal dynamics of numerosity processing, we used multivariate decoding analysis and applied time generalization analysis to determine whether neural representations were transient or sustained over time. Early numerosity-related activity was highly time-specific, with neural patterns most accurately captured when training and testing occurred at the same time points, indicating that initial numerosity encoding is brief and transient. As processing progresses, these representations became more temporally generalized, reflecting a shift toward a stable, enduring numerosity code that could potentially interact with symmetry-related segmentation cues.

Based on several shared characteristics, we previously proposed that the symmetry-induced underestimation illusion may arise through mechanisms similar to those underlying connectedness^45,46^. In our view, symmetry may promote perceptual grouping of elements which in turn leads to a reduction in perceived numerosity. This conclusion is supported by striking parallels between the symmetry induced and the connectedness illusions. First, in both cases, underestimation is stronger at low numerosities^45,50^, likely because objects become harder to segment at higher numerosities.

Second, the magnitude of underestimation scales with the strength of the grouping cue—either additional axes of symmetry or an increased number of connected pairs^40,41,45^. Third, in both illusions, the effect is more pronounced in individuals with lower autistic traits, who may rely less on detail-oriented perceptual strategies^45,51^. Finally, underestimation is enhanced when attentional resources are available and diminished under dual-task conditions, suggesting that both illusions depend on cognitive resources supporting grouping^46,52^. The current study further supports our hypothesis and extends this body of behavioral evidence by highlighting parallels at the neural level. In a combined fMRI and EEG study, Fornaciai and Park^27^ reported that the connectedness effect emerged around 150 ms after stimulus onset and could be decoded from activity patterns in area V3. Similarly, our study demonstrates that the temporal dynamics of the symmetry-driven numerosity underestimation illusion closely mirror those observed for the connectedness illusion. Interestingly, although the relatively late latency of the connectedness effect was initially interpreted as reflecting the contribution of re-entrant feedback signals from higher-order areas, subsequent evidence suggests that such feedback is not necessary for connectedness to bias numerosity perception. Specifically, Fornaciai et al.^53^ investigated the interaction between the connectedness illusion and serial dependence—a perceptual bias induced by recent sensory history—while using backward masking to suppress feedback signals without disrupting feedforward processing. They found that serial dependence was modulated by perceived numerosity, which was reduced by connectedness, rather than by physical numerosity, even when backward masking prevented conscious perception of the connected stimuli. While this does not rule out a role for feedback when stimuli are consciously perceived, these findings suggest that the perceptual organization induced by connectedness may operate automatically through feedforward mechanisms alone. In particular, the authors suggest that it may rely on multiple feedforward sweeps occurring at different times: an initial sweep conveying low–spatial-frequency information, followed by a second sweep that refines perceptual units using high–spatial-frequency signals carried by the connecting lines. Future work should test whether symmetry can similarly bias numerosity perception outside of awareness, as observed for connectedness, or whether the absence of explicit high–spatial-frequency connections precludes the second feedforward sweep. In that case, the late activity observed here would more likely reflect re-entrant feedback processes rather than purely feedforward dynamics. This latter possibility would be more consistent with interpreting the symmetry-induced numerosity underestimation illusion as being mediated by incremental grouping mechanisms that progressively bind items together^46^. According to the incremental grouping theory, perceptual grouping occurs via two processes: base grouping and incremental grouping^54^. Base grouping is rapid, parallel across the visual field, and relies on feedforward activation of neurons tuned to specific features or feature conjunctions (e.g., color, orientation, or their combinations). In contrast, when no neurons are directly tuned to a feature or feature conjunction—such as with symmetry or the combination of numerosity and symmetry—incremental grouping is engaged. This process is slower, capacity-limited, and attention-dependent. During incremental grouping, neurons representing grouped features enhance their activity through recurrent connections, facilitating the co-selection of elements that define the contours of complex shapes, and relying on the spread of attention across elements belonging to the same perceptual object. This view is supported by behavioral evidence showing that reducing attentional resources by means of a concurrent color–orientation conjunction task impairs the detection of symmetry-based groups and diminishes the underestimation effect^46^. However, the involvement of attention does not necessarily imply that the segmentation process relies exclusively on recurrent connections rather than on feedforward sweeps. Indeed, attentional depletion also reduces the connectedness illusion^52^, which, as discussed above, can still operate outside of conscious awareness with no need of feedback signals.

Although our study was not designed to specifically probe the role of attention in symmetry perception or to disentangle feedforward from feedback contributions, a similar debate exists in the symmetry literature. Early research using figure–ground segregation tasks showed that symmetry is detected even under very brief stimulus presentations or by neglect patients, suggesting a pre-attentive processing of symmetry^55,56,57–59^. Yet, although symmetry can be detected even with presentation times of 20 ms^59^, symmetry detection sensitivity improves with longer stimulus durations or with dynamic flicker^60–62^. In addition, studies using multi-color patterns indicate that directing attention to the features of symmetric elements, such as color, enhances symmetry detection, implying a contribution of feature-based attention^63^. These findings suggest that early symmetry detection is driven by rapid feedforward processing, while prolonged viewing, through feedback loops that recruit higher-order attention-dependent regions, facilitates perceptual refinement and greater discrimination precision.

Consistent evidence for this view comes from ERP studies showing that symmetric patterns elicit a relatively late, sustained negative component, the sustained posterior negativity (SPN) which is clearly evident in our VEPs, and thought to represent the neural signature indicating the integration of local visual elements into unified global configurations^64–66^. Although the SPN can arise automatically, even in the absence of sustained stimulation, brief presentation times as short as 20 ms and minimal influence related to response selection^59^, it remains sensitive to top-down contextual priors, such as uncertainty in symmetry-axis orientation^66^, and is modulated by feature-based attention^67^. fMRI studies have also shown that BOLD responses to symmetry are modulated by attention across both dorsal and ventral streams: diverting attention away from symmetric stimuli via a color– or luminance-detection tasks reduces symmetry-related activity^68,69^, whereas explicitly attending to symmetry through a symmetry-detection task enhances these responses^70^.

Interestingly, recent evidence suggests that numerosity is also encoded across both the dorsal and ventral streams^31,32^ and that the signals across the high order areas, at least across the dorsal stream, are modulated by attention^31^. While our EEG results indicate that the interplay between symmetry and numerosity is reflected in occipito-parietal and central electrodes, future research should investigate contributions from both streams with fMRI or MEG to fully characterize the neural substrates underlying symmetry–numerosity interactions.

In summary, the present EEG study demonstrates that numerosity is represented across multiple stages of visual processing: an early stage reflecting raw/physical numerosity, and a later stage that more closely corresponds to perceived numerosity. In both stages, numerosity is encoded independently of other low-level visual features such as dot size and convex hull, consistent with prior fMRI findings^31,32^. The later numerosity-related responses are more sustained, potentially allowing for interactions with symmetry-based cues. Notably, the temporal dynamics of the symmetry-driven numerosity underestimation illusion resemble those reported for the connectedness illusion^27^, suggesting that symmetry-induced underestimation may also emerge at intermediate/high stages of visual processing and is triggered by connectedness-like mechanisms. We propose that this effect arises from later-stage perceptual grouping mechanisms, whereby numerosity is computed from perceptually segmented objects. Supporting this view, the current study shows that numerosity signals interact with symmetry only in a relatively late temporal window, possibly suggesting that attentional and incremental grouping processes may be engaged for symmetry to influence numerosity perception.

## Materials and Methods

### Participants

Twenty-three adult volunteers (4 men, 19 women; mean ± SD age: 24 ± 5) took part in the study. The sample size was based on previous EEG studies investigating similar phenomena, using comparable methodologies^27,59^. Four participants were excluded due to technical issues during EEG recording, resulting in a final sample of 19 participants. All participants had normal or corrected-to-normal vision and were naïve to the purpose of the study. Written informed consent was obtained from all participants, and the study was conducted in accordance with the ethical guidelines of the Commission for Research Ethics at the University of Florence (protocol no. 174, September 23, 2021), and the Declaration of Helsinki.

### Stimuli and experimental procedure

Stimuli were generated using MATLAB (r2016b; MathWorks) and Psychtoolbox routines^71^ and displayed on an AOC Gaming 27G2U/BK monitor (1920×1080 pixels), positioned 57 cm from the observer in a quiet, dark room. Stimuli consisted of arrays of black dots, with varying numerosities: 10, 12, 22 and 24. A red fixation point (0.2° in diameter) remained visible on screen at the center throughout the experiment.

Dots were arranged either symmetrically relative to the vertical and horizontal axes (symmetric condition), or in random positions (random condition). Dot positions were constrained within a an annular region spanning 2-5° of radius from the central fixation (Figure 1). To control for low-level visual features across numerosities, stimulus arrays were generated offline. In half of the trials, convex hull was equated across numerosities, and in the other half, density was equated. The size of individual dots was kept constant at 0.3°, except for numerosity 12, where half of the trials included 0.6° dots to match the overall total surface area of the 24-dot stimuli. Numerosities 10 and 22 were presented only in their random configurations, as they served as perceptual matches for the symmetric 12– and 24-dot stimuli, respectively, based on estimates from previous behavioral studies^45,46^. These combinations resulted in a total of 16 experimental conditions, each repeated 102 times (see Table 1). The order of stimulus presentation was randomized for each participant.

**Table 1.** Experimental conditions included in the study. The table reports all combinations of numerosity (N), spatial arrangement (symmetric; random), non-numerical features (convex hull, density, individual item size / total surface area). Bold numbers indicate stimuli matched for the non-numerical feature specified in the column header. Each condition was repeated 102 times.

| Numerosity (N) | Spatial Arrangement (symm. or rnd.) | Convex hull (deg<sup>2</sup>) | Density (dots/deg<sup>2</sup>) | Item size (size or total surface area matched) |
| --- | --- | --- | --- | --- |
| **10** | *random* | 19 | **0.5** | 0.3° |
|  |  | **56** | 0.2 |  |
| **12** | *symmetry* | 22 | **0.5** | 0.3° |
|  |  | **59** | 0.2 |  |
|  | *random* | 23 | **0.5** |  |
|  |  | **57** | 0.2 |  |
|  | *symmetry* | 22 | **0.5** | 0.6° |
|  |  | **59** | 0.20 |  |
|  | *random* | 23 | **0.5** |  |
|  |  | **57** | 0.2 |  |
| **22** | *random* | 42 | **0.5** | 0.3° |
|  |  | **58** | 0.4 |  |
| **24** | *symmetry* | 44 | **0.5** | 0.3° |
|  |  | **59** | 0.4 |  |
|  | *random* | 46 | **0.5** |  |
|  |  | **58** | 0.4 |  |

Participants were instructed to maintain gaze on the central point. Each stimulus was presented for 150 ms, followed by a variable inter-stimulus interval of 500–700 ms. To maintain attention, in 10% of trials, the dot array was presented in red, and participants compared its numerosity to the immediately preceding black array. They indicated whether the red array appeared more numerous (right arrow) or less numerous (left arrow). These attention trials were excluded from all analyses.

### Electrophysiological Recording and Analysis

EEG activity was recorded continuously throughout the experimental session at a sampling rate of 500 Hz, using a g.Tech g.Nautilus Multi-Purpose system with 30 active electrodes, arranged according to the 10/20 system. Signals were referenced to the right earlobe, and electrode impedance was checked at the beginning of each session, kept below 30 kΩ for all electrodes. A photocell attached to the monitor triggered stimulus onset, ensuring precise synchronization between visual stimulation and EEG recordings.

EEG data were processed in MATLAB (R2013b), using functions from the EEGLAB toolbox^72^, Fieldtrip^73^, and custom scripts. Continous signals were high-pass filtered at 0.1 Hz using a finite impulse response filter with a 0.2 Hz transition bandwidth and a Blackman window. Line noise was attenuated using a notch filter at 49-51 Hz.

The continuous EEG was then segmented into epochs from –150 ms to 600 ms relative to stimulus onset. Baseline correction was applied using the –150 to 0 ms pre-stimulus interval. Epochs were visually inspected, and trials containing artifacts were manually rejected. On average, 79 ± 14 (average ±1 standard deviation) trials per condition were kept for further analyses. Remaining epochs were re-referenced to the common average and averaged separately for each condition. Finally, a low-pass filter at 35 Hz was applied prior to calculating the grand averages.

### Region of interest (ROI)

Consistent with previous studies (Fornaciai & Park, 2017; Park et al., 2016; Bertamini, Rampone et al., 2019; Makin, Wilton et al., 2012) we focused our analyses on occipital and parieto-occipital electrodes, which are known to exhibit the strongest responses to both numerosity^27^ and spatial arrangement of items, such as symmetry^74,59^. The electrodes included in this region of interest (ROI) were O1, O2, PO3, PO4, PO7, PO8, and POz. Analyses on the ROI were performed on the average response across all these electrodes.

### Univariate analysis (VEPs)

We first assessed the effects of numerosity (12 vs. 24), spatial arrangement (random vs. symmetric), and their interaction across all electrodes with a univariate approach. At each time point and electrode, visual evoked potentials (VEPs) were modeled using a linear mixed-effects model, with numerosity (*num*), spatial arrangement (*spat*), and their interaction as fixed effects. To account for the repeated-measures design, random intercepts were included for subjects and for their interactions with the fixed factors. Models were estimated using the Restricted Maximum Likelihood method, following the procedure outlined in Lohse et al^75^. The linear mixed-effect model was specified as:

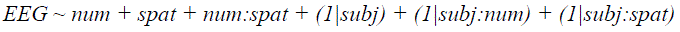

F-statistics were computed using Satterthwaite’s approximation for degrees of freedom. This analysis was then repeated on a restricted ROI, with p-values corrected for multiple comparisons using the False Discovery Rate procedure (FDR, q=0.05).

### Representation similarity analysis (RSA)

Next, we performed representational similarity analysis (RSA)^33,34^, which combined the sensitivity of multivariate approaches with the ability to assess the unique contribution of multiple stimulus dimension to neural activity patterns.

For the parieto-occipital ROI and for each time point, we first computed a neural representational dissimilarity matrix (neural RDM) by calculating the correlation distance (1 – Pearson correlation) between activation patterns for all possible pairs of conditions.

The resulting neural RDMs were entered into a multiple regression analysis with four predictor matrices encoding dissimilarity along key quantitative dimensions of the dot arrays (Euclidean distance): physical numerosity (logarithmic scale), item size, convex hull, and spatial arrangement. This analysis included numerosities 12 and 24, as well as 10 and 22. Previous work indicates that symmetric dot arrays in this range are perceived as approximately 12–14% less numerous than non-symmetric arrays^45^, such that the physical numerosities 10 and 22 in the random condition correspond perceptually to 12 and 24 in the symmetric condition. Analyses were conducted using the *cosmo_target_dsm_corr_measure* function in the CoSMoMVPA toolbox^76^.

Beta weights for each predictor at each time point were tested against zero across subjects using two-tailed one-sample t-tests, with multiple comparisons controlled via FDR (q = 0.05).

Because spatial arrangement influences perceived numerosity, we also investigated whether the neural signal encoded perceived numerosity. The regression analysis was repeated with the physical numerosity regressor replaced by a perceptually adjusted numerosity regressor, derived from prior behavioral estimates (Maldonado Moscoso et al., 2022). Symmetric 12-dot arrays were coded as 10.32 (∼14% underestimation) and symmetric 24-dot arrays as 21.12 (∼12% underestimation). Beta weights for physical versus perceived numerosity were compared using two-tailed paired t-tests, with FDR correction applied (q=0.05).

### Multivariate Pattern Analysis (MVPA)

To complement RSA and capture the contribution of individual electrodes to the pattern of evoked responses as well as additional distributed neural effects over time, we applied multivariate pattern analysis (MVPA) using the MVPA-Light toolbox^77^ to perform both searchlight and temporal generalization analyses^48^.

For classification, we trained a linear discriminant analysis (LDA) classifier using a 5-fold cross-validation procedure, repeated five times with stratification across classes to ensure robust and stable estimates of decoding performance.

The classifier was trained on a subset of EEG data comprising trials from each participant, electrode, and time point (–200 ms to 500 ms relative to stimulus onset). Only trials with numerosities of 12 or 24 dots, presented in either symmetric or random spatial arrangements, were included. Importantly, all stimuli were equated for overall luminance to ensure that decoding performance reflected numerosity and spatial arrangement rather than low-level visual differences.

Two separate classifiers were trained and tested on orthogonal class distinctions, with an equal number of trials per class: (*i*) *numerosity decoding* – the classifier was trained to discriminate between 12 and 24 items, regardless of spatial arrangement; (*ii*) *spatial arrangement decoding* – the classifier was trained to discriminate between random and symmetric arrangements, regardless of numerosity. To assess statistical significance, we employed a cluster-based permutation test (1000 permutations) as implemented in MVPA-Light. Classification accuracies were compared against chance level (50% chance) using a one-tailed Wilcoxon signed-rank test, with cluster correction for multiple comparisons (α = 0.05; cluster statistic = maxsum; cluster criterion = 1.96; minimal connectivity).

To identify the electrodes contributing most to classification performance, we performed a searchlight analysis. Specifically, a classification analysis was conducted independently for each electrode, using time points as features. As in the main MVPA, we employed an LDA classifier with 5-fold cross-validation repeated five times, stratified across classes. Classification accuracy was computed separately for each electrode, providing an estimate of the spatial distribution of decoding performance across the scalp.

For the numerosity classifier, the analysis was restricted to time points between 75–100 ms, whereas for the spatial arrangement classifier we focused on the 140–470 ms window. This design allowed us to probe electrode-specific contributions within the temporal ranges most relevant for each effect. Statistical significance was assessed using the same cluster-based permutation procedure described for the main MVPA, with multiple comparisons correction at α = 0.05.

Finally, we tested the temporal generalization of the classifiers by training and testing them across different time points in the dataset. This approach allows us to determine whether discriminative patterns learned at a given time window remain stable and generalize to other temporal intervals, thereby revealing the temporal dynamics of neural coding. Cross-validation was performed in the same way as in the main MVPA (5-fold, repeated five times, stratified across classes).

## Acknowledgements

This research was funded by the European Union—Next GenerationEU (PRIN 2022, Project RIGHTSTRESS—Tuning arousal for optimal perception, Grant no. 2022CCPJ3J, CUP: B53D23014530001); by the Italian Ministry of Education, University, and Research under the PRIN2017 program (Grant no. 2017XBJN4F—“EnvironMag”).

## References

1. Izard, V., Dehaene-Lambertz, G. & Dehaene, S. Distinct cerebral pathways for object identity and number in human infants. PLoS Biol. 6, e11 (2008).

2. Rugani, R., Regolin, L. & Vallortigara, G. Imprinted numbers: newborn chicks’ sensitivity to number vs. continuous extent of objects they have been reared with. Dev. Sci. 13, 790–797 (2010).

3. Nieder, A. The neuronal code for number. Nat. Rev. Neurosci. 17, 366–382 (2016).

4. Nieder, A. Number Faculty Is Rooted in Our Biological Heritage. Trends Cogn. Sci. 21, 403–404 (2017).

5. Giurfa, M. An Insect’s Sense of Number. Trends Cogn. Sci. 23, 720–722 (2019).

6. Messina, A. et al. Response to change in the number of visual stimuli in zebrafish:A behavioural and molecular study. Sci. Rep. 10, 5769 (2020).

7. Agrillo, C. & Petrazzini, M. E. M. Numerical Competence in Fish. in The Cambridge Handbook of Animal Cognition (eds Kaufman, A. B., Call, J. & Kaufman, J. C.) 580–601 (Cambridge University Press, 2021). doi:10.1017/9781108564113.031.

8. Lorenzi, E., Kobylkov, D. & Vallortigara, G. Is there an innate sense of number in the brain? Cereb. Cortex 35, bhaf004 (2025).

9. Nieder, A. Numerosity coding in the brain: from early visual processing to abstract representations. Cereb. Cortex 35, bhaf180 (2025).

10. Regolin, L. et al. Numerical cognition in birds. Nat. Rev. Psychol. 4, 576–590 (2025).

11. Castaldi, E., Burr, D., Turi, M. & Binda, P. Fast saccadic eye-movements in humans suggest that numerosity perception is automatic and direct. Proc. R. Soc. B Biol. Sci. 287, 20201884 (2020).

12. Castaldi, E., Pomè, A., Cicchini, G. M., Burr, D. & Binda, P. The pupil responds spontaneously to perceived numerosity. Nat. Commun. 12, 5944 (2021).

13. Caponi, C., Castaldi, E., Burr, D. C. & Binda, P. Adaptation to numerosity affects the pupillary light response. Sci. Rep. 14, 6097 (2024).

14. Cicchini, G. M., Anobile, G. & Burr, D. Spontaneous perception of numerosity in humans. Nat. Commun. 7, 12536 (2016).

15. Ferrigno, S., Jara-Ettinger, J., Piantadosi, S. T. & Cantlon, J. F. Universal and uniquely human factors in spontaneous number perception. Nat. Commun. 8, 13968 (2017).

16. Castaldi, E., Mirassou, A., Dehaene, S., Piazza, M. & Eger, E. Asymmetrical interference between number and item size perception provides evidence for a domain specific impairment in dyscalculia. PLOS ONE 13, e0209256 (2018).

17. Burr, D. & Ross, J. A Visual Sense of Number. Curr. Biol. 18, 425–428 (2008).

18. Ross, J. Vision senses number directly. J. Vis. 10, 1–8 (2010).

19. Anobile, G., Cicchini, G. M. & Burr, D. Number As a Primary Perceptual Attribute: A Review. Perception 45, 5–31 (2016).

20. Wagener, L., Loconsole, M., Ditz, H. M. & Nieder, A. Neurons in the Endbrain of Numerically Naive Crows Spontaneously Encode Visual Numerosity. Curr. Biol. 28, 1090–1094.e4 (2018).

21. Ditz, H. M. & Nieder, A. Format-dependent and format-independent representation of sequential and simultaneous numerosity in the crow endbrain. Nat. Commun. 11, 686 (2020).

22. Kutter, E. F., Bostroem, J., Elger, C. E., Mormann, F. & Nieder, A. Single Neurons in the Human Brain Encode Numbers. Neuron 100, 753–761.e4 (2018).

23. Park, J., DeWind, N. K., Woldorff, M. G. & Brannon, E. M. Rapid and Direct Encoding of Numerosity in the Visual Stream. Cereb. Cortex bhv017 (2015) doi:10.1093/cercor/bhv017.

24. Harvey, B. M., Klein, B. P., Petridou, N. & Dumoulin, S. O. Topographic Representation of Numerosity in the Human Parietal Cortex. Science 341, 1123–1126 (2013).

25. Harvey, B. M., Ferri, S. & Orban, G. A. Comparing Parietal Quantity-Processing Mechanisms between Humans and Macaques. Trends Cogn. Sci. 21, 779–793 (2017).

26. Fornaciai, M., Brannon, E. M., Woldorff, M. G. & Park, J. Numerosity processing in early visual cortex. NeuroImage 157, 429–438 (2017).

27. Fornaciai, M. & Park, J. Neural Sensitivity to Numerosity in Early Visual Cortex Is Not Sufficient for the Representation of Numerical Magnitude. J. Cogn. Neurosci. 1–15 (2018) doi:10.1162/jocn_a_01320.

28. DeWind, N. K., Park, J., Woldorff, M. G. & Brannon, E. M. Numerical encoding in early visual cortex. Cortex https://doi.org/10.1016/j.cortex.2018.03.027 (2018) doi:10.1016/j.cortex.2018.03.027.

29. Lasne, G., Piazza, M., Dehaene, S., Kleinschmidt, A. & Eger, E. Discriminability of numerosity-evoked fMRI activity patterns in human intra-parietal cortex reflects behavioral numerical acuity. Cortex Epub ahead of print (2018) doi:10.1016/j.cortex.2018.03.008.

30. DeWind, N. K., Adams, G. K., Platt, M. L. & Brannon, E. M. Modeling the approximate number system to quantify the contribution of visual stimulus features. Cognition 142, 247–265 (2015).

31. Castaldi, E., Piazza, M., Dehaene, S., Vignaud, A. & Eger, E. Attentional amplification of neural codes for number independent of other quantities along the dorsal visual stream. eLife 8, e45160 (2019).

32. Karami, A., Castaldi, E., Eger, E. & Piazza, M. Distinct neural representational geometries of numerosity in early visual and association regions across visual streams. *Commun*. Biol. 8, 1029 (2025).

33. Kriegeskorte, N. Representational similarity analysis – connecting the branches of systems neuroscience. Front. Syst. Neurosci. https://doi.org/10.3389/neuro.06.004.2008 (2008) doi:10.3389/neuro.06.004.2008.

34. Kriegeskorte, N. & Kievit, R. A. Representational geometry: integrating cognition, computation, and the brain. Trends Cogn. Sci. 17, 401–412 (2013).

35. Ginsburg, N. Effect of Item Arrangement on Perceived Numerosity: Randomness vs Regularity. Percept. Mot. Skills 43, 663–668 (1976).

36. Ginsburg, N. & Goldstein, S. R. Measurement of Visual Cluster. Am. J. Psychol. 100, 193 (1987).

37. Im, H. Y., Zhong, S. & Halberda, J. Grouping by proximity and the visual impression of approximate number in random dot arrays. Vision Res. 126, 291–307 (2016).

38. Bertamini, M. Phenomenology, Quantity, and Numerosity. J. Intell. 11, 197 (2023).

39. Franconeri, S. L., Bemis, D. K. & Alvarez, G. A. Number estimation relies on a set of segmented objects. Cognition 113, 1–13 (2009).

40. He, L., Zhang, J., Zhou, T. & Chen, L. Connectedness affects dot numerosity judgment: Implications for configural processing. Psychon. Bull. Rev. 16, 509–517 (2009).

41. He, L., Zhou, K., Zhou, T., He, S. & Chen, L. Topology-defined units in numerosity perception. Proc. Natl. Acad. Sci. 112, E5647–E5655 (2015).

42. Adriano, A., Rinaldi, L. & Girelli, L. Visual illusions as a tool to hijack numerical perception: Disentangling nonsymbolic number from its continuous visual properties. J. Exp. Psychol. Hum. Percept. Perform. 47, 423–441 (2021).

43. Adriano, A. & Ciccione, L. The interplay between spatial and non-spatial grouping cues over approximate number perception. Atten. Percept. Psychophys. 86, 1668–1680 (2024).

44. Apthorp, D. & Bell, J. Symmetry is less than meets the eye. Curr. Biol. 25, R267–R268 (2015).

45. Maldonado Moscoso, P. A., Anobile, G., Burr, D. C., Arrighi, R. & Castaldi, E. Symmetry as a grouping cue for numerosity perception. Sci. Rep. 12, 14418 (2022).

46. Maldonado Moscoso, P. A., Maduli, G., Anobile, G., Arrighi, R. & Castaldi, E. The symmetry-induced numerosity illusion depends on visual attention. Sci. Rep. 13, 12509 (2023).

47. Anobile, G., Arrighi, R., Castaldi, E. & Burr, D. C. A Sensorimotor Numerosity System. Trends Cogn. Sci. 25, 24–36 (2021).

48. King, J.-R. & Dehaene, S. Characterizing the dynamics of mental representations: the temporal generalization method. Trends Cogn. Sci. 18, 203–210 (2014).

49. Karami, A., Castaldi, E., Eger, E., Hebart, M. & Piazza, M. Numerosity Is Directly Sensed and Dynamically Transformed in the Human Brain: Evidence from MEG-MRI Fusion. Preprint at 10.1101/2025.11.15.687894 (2025).

50. Anobile, G., Cicchini, G. M., Pomè, A. & Burr, D. C. Connecting visual objects reduces perceived numerosity and density for sparse but not dense patterns. J. Numer. Cogn. 3, 133–146 (2017).

51. Pomè, A., Caponi, C. & Burr, D. C. Grouping-Induced Numerosity Biases Vary with Autistic-Like Personality Traits. J. Autism Dev. Disord. 52, 1326–1333 (2022).

52. Pomè, A., Caponi, C. & Burr, D. C. The Grouping-Induced Numerosity Illusion Is Attention-Dependent. Front. Hum. Neurosci. 15, 745188 (2021).

53. Fornaciai, M. & Park, J. Disentangling feedforward versus feedback processing in numerosity representation. Cortex 135, 255–267 (2021).

54. Roelfsema, P. R. & Houtkamp, R. Incremental grouping of image elements in vision. Atten. Percept. Psychophys. 73, 2542–2572 (2011).

55. Driver, J., Baylis, G. C. & Rafal, R. D. Preserved figure-ground segregation and symmetry perception in visual neglect. Nature 360, 73–75 (1992).

56. Locher, P. J. & Wagemans, J. Effects of Element Type and Spatial Grouping on Symmetry Detection. Perception 22, 565–587 (1993).

57. Treder, M. S. Behind the Looking-Glass: A Review on Human Symmetry Perception. Symmetry 2, 1510–1543 (2010).

58. Makin, A. D. J., Roccato, M., Karakashevska, E., Tyson-Carr, J. & Bertamini, M. Symmetry Perception and Psychedelic Experience. Symmetry 15, 1340 (2023).

59. Contemori, G., et al. Sustained Posterior Negativity (SPN) Elicited by Brief (20ms) Symmetrical Stimuli. Frontiers in Psychology vol. (In press) (2025).

60. Tyler, C. W., Hardage, L. & Miller, R. T. Multiple mechanisms for the detection of mirror symmetry. Spat. Vis. 9, 79–100 (1995).

61. Niimi, R., Watanabe, K. & Yokosawa, K. The dynamic-stimulus advantage of visual symmetry perception. Psychol. Res. 72, 567–579 (2008).

62. Sharman, R. J. & Gheorghiu, E. The role of motion and number of element locations in mirror symmetry perception. Sci. Rep. 7, 45679 (2017).

63. Gheorghiu, E., Kingdom, F. A. A., Remkes, A., Li, H.-C. O. & Rainville, S. The role of color and attention-to-color in mirror-symmetry perception. Sci. Rep. 6, 29287 (2016).

64. Norcia, A. M., Candy, T. R., Pettet, M. W., Vildavski, V. Y. & Tyler, C. W. Temporal dynamics of the human response to symmetry. J. Vis. 2, 1–1 (2002).

65. Höfel, L. & Jacobsen, T. Electrophysiological indices of processing aesthetics: Spontaneous or intentional processes? Int. J. Psychophysiol. 65, 20–31 (2007).

66. Makin, A. D. J., Buckley, N., Austin, E. & Bertamini, M. When does perceptual organization happen? Cortex 174, 70–92 (2024).

67. Bertamini, M., Rampone, G., Tyson-Carr, J. & Makin, A. D. J. The response to symmetry in extrastriate areas and its time course are modulated by selective attention. Vision Res. 177, 68–75 (2020).

68. Sasaki, Y., Vanduffel, W., Knutsen, T., Tyler, C. & Tootell, R. Symmetry activates extrastriate visual cortex in human and nonhuman primates. Proc. Natl. Acad. Sci. 102, 3159–3163 (2005).

69. Zamboni, E., Makin, A. D. J., Bertamini, M. & Morland, A. B. The role of task on the human brain’s responses to, and representation of, visual regularity defined by reflection and rotation. NeuroImage 297, 120760 (2024).

70. Keefe, B. D. et al. Emergence of symmetry selectivity in the visual areas of the human brain: fMRI responses to symmetry presented in both frontoparallel and slanted planes. Hum. Brain Mapp. 39, 3813–3826 (2018).

71. Brainard, D. H. The Psychophysics Toolbox. Spat. Vis. 10, 433–436 (1997).

72. Delorme, A. & Makeig, S. EEGLAB: an open source toolbox for analysis of single-trial EEG dynamics including independent component analysis. J. Neurosci. Methods 134, 9–21 (2004).

73. Oostenveld, R., Fries, P., Maris, E. & Schoffelen, J.-M. FieldTrip: Open Source Software for Advanced Analysis of MEG, EEG, and Invasive Electrophysiological Data. Comput. Intell. Neurosci. 2011, 1–9 (2011).

74. Makin, A. D. J., Wilton, M. M., Pecchinenda, A. & Bertamini, M. Symmetry perception and affective responses: A combined EEG/EMG study. Neuropsychologia 50, 3250–3261 (2012).

75. Lohse, K. R., Kozlowski, A. J. & Strube, M. J. Model Specification in Mixed-Effects Models: A Focus on Random Effects. https://doi.org/10.48550/arXiv.2209.14349 (2022) doi:10.48550/arXiv.2209.14349.

76. Oosterhof, N. N., Connolly, A. C. & Haxby, J. V. CoSMoMVPA: Multi-Modal Multivariate Pattern Analysis of Neuroimaging Data in Matlab/GNU Octave. *Front*. Neuroinformatics 10, (2016).

77. Treder, M. S. MVPA-Light: A Classification and Regression Toolbox for Multi-Dimensional Data. Front. Neurosci. 14, 289 (2020).

